# LRH-1 NUTRIGENOMICS: The Provision of Lauric Acid Results in the Endogenous Production of the Liver Receptor Homolog-1 Ligand, Dilauroylphosphatidylcholine, and LRH-1 Transactivation

**DOI:** 10.1101/2021.02.01.429240

**Authors:** KC Klatt, S. Zhang, OV Malysheva, Z. Sun, B. Dong, JT. Brenna, DD. Moore, MS. Roberson, MA Caudill

## Abstract

**Background:** The unusual phosphatidylcholine species, dilauroylphosphatidylcholine (DLPC), has been reported to bind and activate the orphan nuclear receptor, liver receptor homolog-1 (LRH-1). To date, DLPC has not been reported endogenously in metabolomic databases.

**Objective:** Herein, we test the hypothesis that the provision of the acyl constituent of DLPC, lauric acid (C12:0), a saturated fatty acid rich in tropical oils such as coconut oil, will 1) result in endogenous DLPC production and 2) enhance LRH-1 transcriptional activity.

**Methods:** We measured DLPC following provision of C12:0 to HepG2 cells, C57/BL6J mice, and to healthy human participants in an acute, randomized, controlled cross-over trial. LRH-1^fl/fl^ and LRH-1^fl/fl^ Albumin-Cre mice were used in *ex vivo* and *in vivo* approaches. to assess the impact of C12:0 on LRH-1 target gene expression. 1-^13^C-lauric acid and methyl-d9-choline were used to assess DLPC production dynamics.

**Results:** DLPC was not observed in any C12:0-free approach. Provision of C12:0 in the culture media or to C57/BL6J mice resulted in the rapid production of DLPC, including DLPC’s presence in multiple LRH-1 expressing tissues. Coconut oil-fed human participants exhibited DLPC in postprandial serum samples. *Ex vivo* and i*n vivo* C12:0 provision resulted in increased mRNA expression of LRH-1 target genes, an effect that was not observed in hepatic knockout mice. Methyl-d9-choline administration revealed a complex reliance on CDP-choline-derived DLPC.

**Conclusion:** C12:0 provision results in endogenous production of the LRH-1 ligand, DLPC, and LRH-1 transcriptional activation phenotypes. Our findings highlight pleiotropic effects of lauric acid, a common hypercholesterolemic dietary saturated fatty acid, secondary to LRH-1 agonism.

## INTRODUCTION

Nuclear receptors (NRs) are a superfamily of ligand-activated transcription factors that regulate diverse aspects of mammalian physiology^1^. NRs are deemed either orphaned or adopted based on the identification of an endogenous ligand. Ligand binding to the ligand binding domain (LBD) acts as an allosteric regulator, initiating a series of biophysical events that result in direct regulation of specific target gene transcription in tissue- and cell type-specific manners. Liver receptor homolog-1 (LRH-1; NR5A2) is an orphan nuclear receptor of the NR5A subfamily that is expressed primarily in tissues of endodermal origin, including the liver, intestine, exocrine pancreas, ovary, and pre-adipocytes^2–4^. Several regulatory roles have been described for LRH-1, including the maintenance of pluripotency, metabolism of cholesterol into diverse, tissue-specific products (i.e. bile acids, progesterone, glucocorticoids) and recently as a major regulator of methyl metabolism^5^, making LRH-1 signaling an attractive target for both therapeutic activation and inhibition.

Investigations into the structure of LRH-1 have revealed an LBD occupied by several phospholipid classes; this phospholipid binding appears promiscuous and the ability of phospholipids to result in altered transcriptional activity under physiological conditions remains largely unexplored^6–9^. One report^10^ identified an ‘uncommon’ phosphatidylcholine (PC), dilauroylphopshatidylcholine (DLPC) as a potent agonist ligand of human and mouse LRH-1, able to bind the LRH-1 LBD and induce LRH-1-related transcriptional phenotypes. Despite convincing evidence that DLPC is indeed a LRH-1 ligand^11,12 13^, DLPC has yet to be catalogued in the Human Metabolomics Database or the Mouse Multiple Tissue Metabolomics Database^14^, ^15^.

DLPC is a PC species with dodecanoic/lauric (C12:0) acid esterified at the *sn*-1 and *sn*-2 positions of the glycerol backbone (commonly termed PC 12:0/12:0), an uncommon structure for PC species which typically contain a mix of saturated and unsaturated fats at the sn-1 and sn-2 respectively. While choline, methyl donors and fatty acids required to produce PCs endogenously are common in cell culture media and mammalian diets, C12:0 is typically not abundant. C12:0 is either absent or a trace fatty acid in cell culture media supplied with fetal bovine serum. Considered one of the three primary, dietary hypercholesterolemic saturated fats, C12:0 is only found in very low quantities in common foods, is absent from common chow diets fed to laboratory rodents, and is low in the diets of most humans consuming western-style diets (<1% on kilocalorie intake on average in NHANES; **Supplementary Figure 1**). Endogenous production of C12:0 may occur as an extremely minor product of fatty acid synthesis or as an intermediate in fatty acid oxidation, but it is typically found at very low levels in cultured cells and likely does not exist in exchangeable pools available available PC synthesis^16,17^. It may therefore be considered unsurprising that DLPC has yet to be reported *in vitro* or *in vivo*. The absence of DLPC from physiological systems may additionally be compounded by limited incorporation into membrane phospholipids, suggesting phospholipids containing C12:0 should be labile and unlikely to be detected in commonly available fasted blood samples^18 19^.

While C12:0 intake is low in most laboratory animal diets and typical human western-style diets, it may be supplemented directly in the culture media/diet, or by including tropical oils, such as palm kernel oil and coconut oil, which provide rich sources of C12:0^20–22^ due to the plant’s expression of medium chain acyl-ACP thioesterases. C12:0 is also relatively abundant in murine and human milk, constituting about 5-10% of total fatty acids^23^. Traditional diets of rodent and human populations in Pacific Islands subsisting on coconut rich products can have lauric acid constitutes up >20% of total caloric intake, reflected in high levels of lauric acid in adipose tissue stores^21,24^. High intakes of C12:0-containing products in traditional equatorial diets despite C12:0’s classification as a cholesterol-raising saturated fatty acid have led to substantial debate about its impact on cardiometabolic health. Despite increases in risk factors, particularly LDL, preclinical rodent models employing coconut oil have failed to observe expected atherogenic^25,26^ and diabetogenic^27^ phenotypes. Both clinical trial^28^ and observational epidemiological investigations^29,30^ have suggested beneficial cardiometabolic effects of C12:0 relative to other saturated fatty acids. Such observations are consistent with pleiotropic effects of C12:0, though few have been described to date, apart from C12:0’s relative high oxidation rate *in vivo* ^31^ and ability to activate toll-like receptors at high concentrations *in vitro* ^32–34^.

Herein, we utilize *in vitro*, *ex vivo*, wild type and transgenic murine models, as well as a single blind, acute, randomized controlled cross-over feeding trial in humans to test the hypothesis that the provision of C12:0 in the available nutriture will result in the endogenous production of DLPC, as well as LRH-1 activation phenotypes. Our results support the idea that the lack of endogenous DLPC is due to the limited C12:0 acyl substrate for PC synthesis and suggest that LRH-1 transactivation could contribute to its previously observed pleiotropic effects.

## METHODS

### In Vitro

HepG2 cells (ATCC^®^ HB-8065™) were cultured in Dulbecco’s Modified Eagle Medium (DMEM) containing 10% Fetal Bovine Serum. All experiments with HepG2 cells were performed prior to passage 5. Cell culture medium was supplemented with free fatty acids (NuChek Prep) bound to albumin (USBiological; CAS # 9048-46-8) in Dulbecco’s Phosphate Buffered Saline (DPBS; 1%) in 10 cm^3^ dishes when cells reached 75% confluency. Control treatments received an equivalent volume of fatty acid-free bovine serum albumin-DPBS solution (BSA). Fatty acids and their respective concentrations are indicated in the figure legends. For pulse-chase experiments, C12:0 (100μM) or control was provided for 4 hours. The media was subsequently removed, the cells were washed and replenished with media containing 10% FBS but no supplemental C12:0; cell lysates were progressively harvested at the indicated time points for LC/MS analysis.

### Ex Vivo

Hepatocyte-specific LRH-1 knockout mice were generated via breeding LRH-1^fl/fl^ mice with Albumin-Cre mice as previously described ^35^. 12-week old hepatic LRH-1 wild type (Cre^−^; hLRH1-WT; n=2) and knockout (Cre^+^; hLRH1-KO; n=2) mice underwent primary hepatocyte isolation, as previously described ^35^. Briefly, mice were euthanized with isoflurane and perfused via inferior vena cava with Earle’s balanced salt solution containing 5mM EGTA followed by 100U/mL collagenase supplemented Hank’s balanced salt solution. The liver was removed, massaged to dissociate hepatocytes, filtered, washed and layered onto Percoll for live cell isolation. Following isolation, cells were plated in 5% FBS-containing Williams E media, allowed to rest for 12-16 hours, and subjected to 8 hours of treatment with 100μM of the indicated BSA-bound free fatty acids or volume equivalent of vehicle (BSA). Following treatment, media was aspirated, cells washed, collected in Trizol and snap-frozen for determination of LRH-1 target gene expression via qRT-PCR.

### Animal Dietary Experiments

All animal protocols and procedures used in this study were approved by the Institutional Animal Care and Use Committees at Cornell University (wild -type experiments) or Baylor College of Medicine (transgenic experiments), and were conducted in accordance with the Guide for the Care and Use of Laboratory Animals. C57/BL6J mice were purchased from Jackson Laboratories and housed in microisolator cages in an environmentally controlled room (22-25°C and 70% humidity with a 12-hour light-dark cycle). Prior to the start of experimental diets, all mice consumed chow diets (Teklad 2018).

### Acute, Wild-Type Feeding

#### Lauric Acid

To assess the effects of C12:0 provision on acute production of DLPC, male and female mice (n=3 per group) were gavaged with test oil mixtures, and sacrificed for serum and tissue collection after 1.5 hours. Duodenum (1.5cm) and serum were collected following CO_2_ euthanasia. Fatty acids, oil mixtures, and their doses are as stated in figure legends. A single mouse was gavaged with 125mg of 1-^13^C-lauric acid in order to confirm direct incorporation of exogenous C12:0 into DLPC.

### De Novo Lipogenesis

To confirm the inability of endogenous C12:0 pools to facilitate DLPC production, C57/BL6 male and female mice were fed high sucrose, 0% fat diets (Envigo; TD.03314) for 1, 3, 5, 7 days to stimulate de novo lipogenesis, as previously described ^36^, sacrificed, and metabolic tissues (intestine, liver, pancreas, kidney) were collected for LC/MS analysis.

### Chronic Feeding

To assess the effects of habitual C12:0 consumption on DLPC production, 12 8-week old C57/BL6J mice per group (6 male, 6 female) were randomly assigned to consume chow, olive oil, or coconut oil based high fat diets (60% kcal from fat; 18% kcal from C12:0) for 1 month, simulating traditional C12:0 intakes in Polynesian atolls. Diets were based on chow (Teklad 2018) supplemented with no added fat or with refined oils and vitamin (Dyets Inc., AIN-93VX Vitamin Mix, 310025)/mineral (Dyets Inc., AIN-93G MX Mineral Mix, 210025) mixtures (see **Supplementary Table 1** for recipe). Mice were fasted overnight (~14 hours) and divided into 2 groups: fasted, for immediate sacrifice, or postprandial, who underwent gavage with 50μL of soybean oil (chow), refined olive oil (olive), or refined coconut oil (coconut). The following tissues were collected and snap frozen in liquid nitrogen for LC-MS/MS analysis: serum (post-mortem heart stick), erythrocytes, duodenum (1.5 cm), whole liver, splenic side pancreas, kidney, hindlimb skeletal muscle (rectus femoris, vastus lateralis), whole brain, testis, and ovaries.

### Acute, Transgenic Feeding

At 10-12 weeks of age, male Cre^+^ hepatic LRH-1 knockout mice (hLRH1-KO) and Cre^−^ hepatic LRH-1 wild-type (hLRH1-WT) mice were randomly assigned to 10 days of feeding of custom purified diets (Research Diets, Inc, New Brunswick, NJ) with 21% kcal from fat, and either 10%kcal from C12:0 or an isocaloric comparator, C16:0 (palmitic acid); diet composition can be found in **Supplementary Table 3**. Three separate feeding cohorts utilizing sibling-matched hLRH1-KO and hLRH1-WT mice were undertaken to achieve the final sample size, as follows: hLRH1-WT-C12:0, n=7; hLRH1-WT-C16:0, n=7; hLRH1-KO-C12:0, n=7; hLRH1-KO-C16:0, n=9. Given LRH-1’s feeding-related circadian rhythm ^4,37^, mice were sacrificed after a brief (2 hour) fast in the morning. Liver tissue was harvested and snap-frozen for RNA isolation and qRT-PCR.

### Deuterium-labeled Choline Administration

To assess the contribution of the CDP-choline and PEMT pathway to in vivo production of DLPC, the stable isotope, methyl-d9-choline (Cambridge Isotopes Laboratory, Andover, MA, USA), was administered to 2 female C57/BL6J mice consumed 45% high fat diet (10% C12:0) and drinking water containing 4mM d9-choline for 8 weeks. Mice were sacrificed ad libitum and tissues were harvested for labeled and unlabeled DLPC analysis (see LC-MS methods).

### Acute Randomized, Single-Blind, Cross-Over Feeding Trial in Human Subjects

To assess the impact of C12:0 consumption on DLPC production in humans, we undertook an acute, randomized, single-blind, cross-over feeding trial in 10 healthy, human subject participants (ClinicalTrials.gov NCT034816080). A CONSORT diagram and participant characteristics can be found in **Supplementary Figure 2** and **Supplementary Table 3**, respectively. Participants arrived fasted (~10h) at the Francis A. Johnston and Charlotte M. Young Human Metabolic Research Unit (HMRU) at Cornell University. Participants were randomly assigned to consume a breakfast shake containing either 27.2g of refined coconut (Nutiva Organic, Neutral Tasting) or refined olive oil (Filippo Berio). The unit of randomization was the order in which the study test oils were consumed. Breakfast shakes were prepared using gram scales in the metabolic kitchen at the HMRU. Participants provided a fasted blood draw at each visit, followed by additional blood draws 2, 4 and 6 hours postprandially. Whole blood was allowed to clot for 30 minutes at room temperature and centrifuged at 2000 x g for 15 minutes at 4°C to obtain serum and stored at −80°C. Participants were instructed to not consume additional food prior to their last blood draw; participants were allowed to consume water, and black coffee/tea *ad libitum*. Participants completed the second study visit following at least a 1 week washout period.

### Liquid Chromatography-Mass Spectrometry Analysis of Phosphatidylcholines

Total lipid extraction from serum and other tissues was undertaken using an adaptation of the method of Koc et al ^38^. Briefly, 0.02-0.05 g of solid tissue or 150 μL of serum homogenized in 400μL 2:1 methanol:chloroform with 5nM/L PC 9:0-9:0 (Avanti Polar Lipids, Alabaster, Alabama) as an internal standard. The samples were incubated overnight (18 hrs minimum) at −20°C. Homogenates were centrifuged at 5000 rpm for 5 min and upper fractions were transferred into new tubes. To the remaining of the solid residue, 250μL of 2:1:0.8 (methanol:chloroform:water) was added. After vortexing and centrifugation (5 min 14000 rpm), the upper fraction was combined with the previously collected fraction. To separate water-soluble compounds from lipid-soluble, 100μL of chloroform and 100μL of water were added to the tubes. Following centrifugation, the lower layer was collected and dried in Savant SpeedVac. Extracts were reconstituted in 100μL of 6:1 (methanol:chloroform).

The extract (5μL) was injected on the UPLC-MS system containing an UltiMate 3000 Dionex Autosampler and Pump, and Q-Exactive MS with ESI probe (Thermo Fisher Scientific, Waltham, MA). Total PC was separated from sphingomyelin (SM) and LPC using a Syncronis Sillica column (2.1 x 150 mm, 5um; Thermo) and mobile phase containing buffer A (acetonitrile 400mL, water 127mL, EtOH 68mL, 3mL 1M ammonium acetate, 2mL glacial Acetic acid) and buffer B (acetonitrile 250mL, water 250mL, EtOH 42mL, 13.5mL 1M ammonium acetate, 9mL glacial acetic acid) with gradient elution at 0.4mL/min: from 0 to 3 min isocratic mix with 5% buffer B, from 3 to 10 linear gradient from 5 to 30% buffer B; from 10 to 14 min linear gradient from 30 to 60% buffer B; from 14 to 16 min - 100% buffer B; held for 1 min at 100% buffer B, then from 17-19 min linear gradient from 100 to 5% buffer B; from 19 to 21 min column was equlibrated to starting condition (5% buffer B). Samples were kept at 5°C; the column’s temperature was held at 30°C. The MS was operated in positive (full scan only) and negative (MS2) modes with the following settings: sheath gas pressure was set at 40; auxillary gas 10; spray voltage 5kV; capillary temperature 350°C; S-lens RF level 50; heater temperature 30C. MS2 mode was used for detection and quantification of DLPC, where resolution was set at 35,000; AGC target 2e4; isolation window 1 m/z; HCE 20. The inclusion list contained m/z 680.45 which represented a DLPC anionized adduct [M+OAc]- and m/z 596.33 for PC9:0-9:0. For quantification, the sn1/2 RCOO- ion, m/z 199.17, was used for DLPC and m/z 157.1 was used for PC9:0-9:0. The anionic loss of CH_3_, m/z 606.39, was used as DLPC qualifier. Full scan mode (AIF) was used to quantify total PC. Resolution was set at 35,000; AGC target 3e6; scan range 80-950; HCE 20. For quantification of total PC, a m/z of 184.1 was used. Quantification was performed using XCalibur software (Thermo Fisher Scientific).

### Quantitative Real Time Polymerase Chain Reaction (qRT-PCR)

To assess the ability of lauric-acid induced DLPC production to agonize LRH-1, we measured the relative abundance of the downstream LRH-1 transcripts, *CYP8B1* and *GNMT* using both primary hepatocytes and whole liver tissue obtained from hLRH1-WT and hLRH1-KO. RNA was extracted from primary hepatocytes treated with indicated fatty acids and liver tissue derived from LRH-1^fl/fl^ Alb-Cre feeding studies with Trizol reagent (Invitrogen™) using the manufacturer’s protocol. RNA concentration and quality were assessed using a NanoDrop ND-1000 spectrophotometer (ThermoFisher Scientific). cDNA was generated using a high capacity cDNA reverse transcription kit (Applied Biosystems, Thermo Fisher Scientific) and an Eppendorf 5331 Mastercycler. Quantitative PCR was performed with a Roche LightCycler 480 II using SYBR green supermix reagents (Bio-Rad Laboratories). A panel of reference genes was utilized to determine the most stably expressed reference target (NormFinder, Aarhus University Hospital, Denmark). Forward and reverse primer sequences are shown in **Supplementary Table 2**. Primer efficiencies were calculated following amplification of a standard curve. Melting curves were included at the end of amplification cycles to validate specificity of the PCR product. The ∆,∆Ct method was used to calculate fold changes.

### Statistical Analysis

All statistical analyses reported for DLPC production are within treatment analyses, owing to the lack of DLPC observed outside of contexts where C12:0 was provided. Repeated measures ANOVA were used to determine the significant effects of C12:0 on DLPC production across a time course in cultured cells, and in post-prandial blood samples from humans fed coconut oil diets. 1-way ANOVAs were performed to compare the effect of different C12:0 doses on DLPC production in cultured cells. 2-way ANOVA with Tukey’s post-hoc test was used to assess the impact of fatty acid provision on LRH-1-dependent gene expression.

## RESULTS

### C12:0 Provision Results in Rapid Production of DLPC *In Vitro*

Consistent with the reported lack of DLPC in cultured cells, we observed no DLPC production in any cell lysates from any cells cultured without exogenously provided C12:0. Provision of exogenous C12:0 to HepG2 cells resulted in endogenous DLPC production across a time course (100 μM dosage) and across a range of doses (4 hour treatments) (**Figure 1A;1B**). Consistent with previous data showing that C12:0 exhibits low incorporation into membranes, and that regulation of nuclear receptor signaling occurs by degradation of endogenous ligands, we observed a very short half-life of exogenously added DLPC, estimated to be ~47 min (**Figure 1C**). We also observed DLPC production with 4 hour C12:0 treatment across several diverse cell types, including HeLa, JEG3, C2C12 and Caco-2 cells, suggesting that C12:0 is readily incorporated into fatty acid pools available for incorporation into diacylglycerols and subsequent phosphatidylcholine synthesis across a range of cell types (Supp x and data not shown). Collectively, these findings suggest that C12:0 provision results in DLPC production in cultured cells.

**Figure 1:**
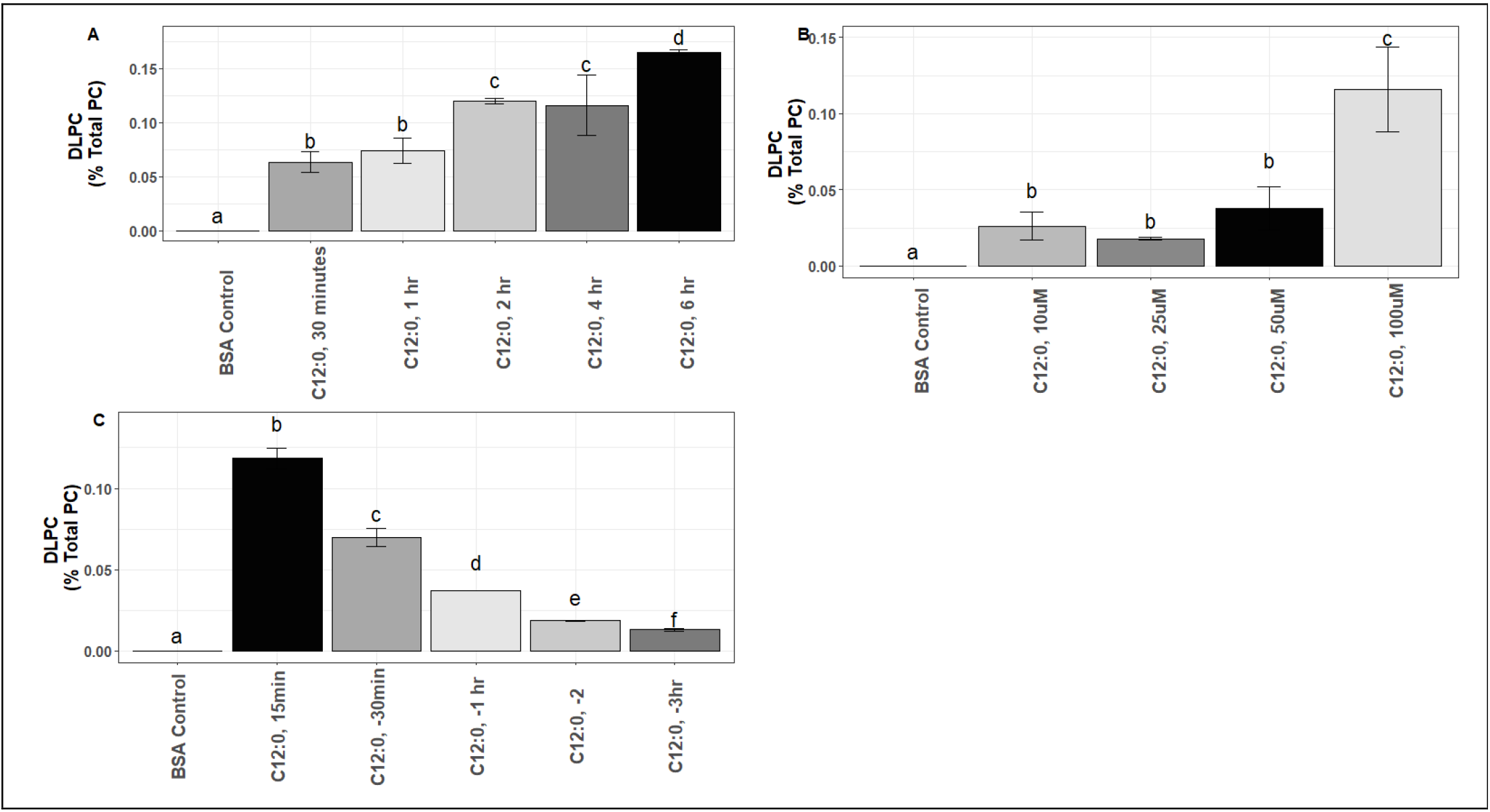
C12:0 Provision Results in Rapid Production of DLPC In Vitro. A) HepG2 cells were cultured across a time course with 100μM of lauric acid. B) HepG2 cells were cultured with varying concentrations (0μM to 100μM) of lauric acid for 4 hours. C) HepG2 cells were cultured with 100μM of lauric acid for 4 hours, washed and replaced with unsupplemented media and harvested at serial time points.

### C12:0 Provision Results in Rapid Intestinal Production of DLPC and Exposure to LRH-1 Expressing Tissues *In Vivo*

We observed rapid production of DLPC in the duodenum of male and female mice following acute (1.5h) oral gavage of olive oil supplemented with free C12:0 and coconut oil, a dietary source of C12:0 (**Figure 2A**). Supplementation ^13^C-C12:0 also resulted in direct incorporation of ^13^C-C12:0 into DLPC (**Figure 2A**). As predicted from the absence of endogenous C12:0in exchangeable pools available for PC biosynthesis, only DLPC molecules containing two ^13^C-C12:0s detected. DLPC was also detected at low concentrations in postprandial serum (**Figure 2B**). Chronic feeding (1 month) of high fat coconut oil diets (60% kcal; 18% lauric acid) resulted in low but detectable presence of DLPC in multiple organs in addition to the intestine, including kidney, liver, skeletal muscle and ovary (**Figure 2C**). DLPC was not observed in fasting serum samples, washed red blood cells, brain, or testes. These concentrations of DLPC were measured after overnight fasting and likely underestimate tissue exposure to some degree, based on the rapid degradation of DLPC observed *in vitro*.

**Figure 2:**
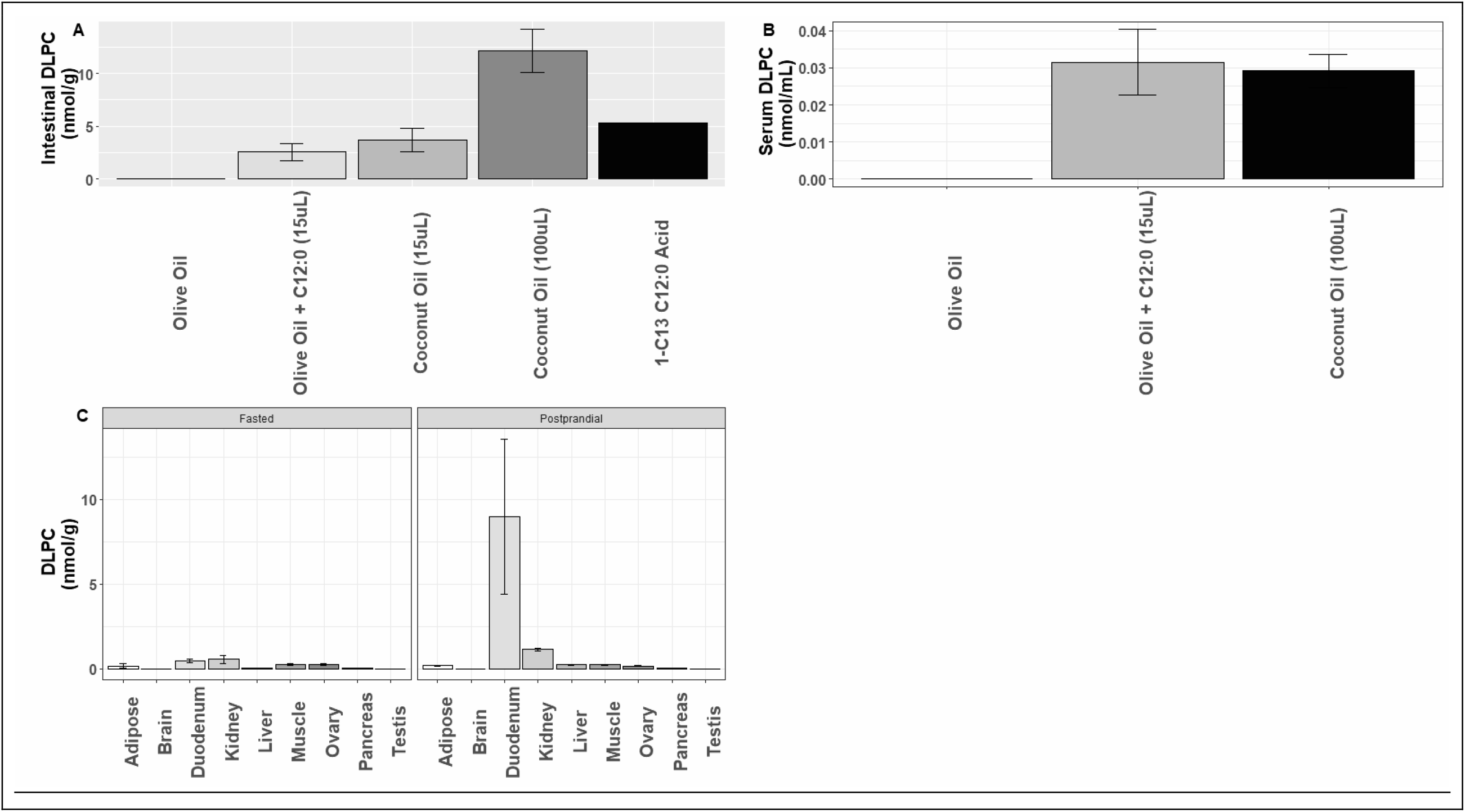
C12:0 Provision Results in Rapid Intestinal Production of DLPC and Exposure to Broad LRH-1 Expressing Tissues In Vivo. A) Acute, oral gavage (n=3) of C12 containing oils and duodenal DLPC production (~1.5hr post-gavage). B) Appearance of DLPC in postprandial serum. C) Tissue Atlas of DLPC Production in both Fasted (14 hour) and Postprandial (14 hour fast + 3 hour gavage) states (n=3/group).

### De Novo Lipogenic Diets Are Insufficient to Induce Endogenous DLPC Production

To confirm the presumption that de novo lipogenesis is not sufficient to generate a physiological pool of C12:0 available for DLPC production, C57/BL6J male and female mice (n=3 per sex) were fed 0% fat diets over a time course (1, 3, 5, 7 d); feeding a 0% fat, high sucrose diet is a model of enhanced *de novo* lipogenesis, previously shown to generate a fatty acid synthase-derived PC species that acts as an endogenous ligand for the peroxisome proliferator activated receptor alpha^36^. We did not detect DLPC in major metabolic tissues (intestine, liver, pancreas, kidney) following this lipogenic diet. These results support our hypothesis that DLPC production requires exogenous provision of C12:0 in the diet/culture media.

### Deuterium-Labeled Choline Reveals Intestinal CDP-Choline-Derived DLPC with Broad Extra-Intestinal Remodeling of PC Species to Facilitate Endogenous DLPC Production

Production of di-saturated PC species has been reported with specific tissues exhibiting the capacity to produce high quantities, such as the lung’s production of the surfactant, dipalmitoylphosphatidylcholine. It remains unclear whether intestinal production of DLPC is the predominant source of DLPC for extra-intestinal tissues, or whether other tissue types are able to produce DLPC when presented with C12:0. To begin to address this, we provided female mice d9-methyl-choline in their drinking water for 4 weeks; this deuterated form of choline has been used to assess production of PC species through the CDP-choline vs the Phosphatidylethanolamine N-Methyltransferase (PEMT) pathway of PC synthesis^39,40^. PC produced through the CDP-choline pathway retains all 9 deuterated methyl groups (d9-PC), whereas PEMT-derived-PC predominantly exhibits a d3-label (d3-PC), following incorporation of d9-choline label into the methyl pool. d3-PC also results from the CDP-choline pathway, following phospholipase D activity on PEMT-derived-PC, liberating a d3-choline moiety to be utilized by tissues via the CDP-choline pathway. In the *ad libitum* state, d9-methyl-choline supplementation resulted in comparable amounts of total d9- and d3-PC in the intestine, indicating contributions of both pathways (**Table 1**). In contrast, the striking 100x predominance of intestinal d9-DLPC relative to d3-DPLC indicates dependence on the CDP-choline pathway, which is consistent with the reported lack of detectable intestinal PEMT expression/activity^41^. Unlike the intestine, and contrary to the notion that the intestine supplies DLPC for the mammalian body, plasma enrichments exhibited near equivalent ratios of d3- and d9-DLPC. Notably, the liver, an organ with very high expression of PEMT, showed low levels of d9-DLPC but undetectable levels of d3-DLPC, despite a greater enrichment of d3-PC relative to d9-PC. Collectively, the labeling scheme across these three compartments support a model whereby the CDP-choline pathway is critical for producing DLPC, either through direct condensation of CDP-choline with dlauroylglycerol, or remodeling of CDP-choline-derived PCs. The plasma compartment’s enrichment with d3-DLPC likely results from recycling of the d3-choline moiety from d3-PC through the CDP-choline pathway and is broadly indicative of the extra-intestinal capacity to produce DLPC.

**Table 1:** d9-Choline Provision Reveals CDP-Choline-Dependent, Dynamic DLPC Synthesis from C12:0.

|  | <u>% Total PC Enrichment</u> |  |  |
| --- | --- | --- | --- |
| **Tissue** | <u>Intestine</u> | <u>Serum</u> | <u>Liver</u> |
| **d9-DLPC** | 0.032 | 0.0072 | 0.00030 |
| **d9-PC** | 10.95 | 8.53 | 7.55 |
| **d3-DLPC** | 0.00033 | 0.0075 | Undetectable |
| **d3-PC** | 12.04 | 16.77 | 13.31 |

### C12:0 Provision Results in the Appearance of DLPC in Postprandial Serum of Coconut Oil-Fed Healthy Human Subjects

Healthy human participants (n=10) participated in an acute, randomized, controlled cross-over feeding trial to assess the impact of a C12:0 containing breakfast smoothie (prepared with 11.38g of C12:0 in the form of coconut oil) relative to a C12:0-free control (0g; provided as olive oil). Consistent with DLPC being absent in the Human Metabolomics Database, we did not observe DLPC in any fasting plasma samples from either study visit. We additionally did not observe DLPC in post-prandial serum following consumption of breakfast shakes prepared with olive oil. As hypothesized, we did observe DLPC in postprandial serum following consumption of breakfast shakes prepared with coconut oil (**Figure 3**). Consistent with our results in acute animal gavage experiments, DLPC appeared in serum 2 h postprandially and further increased at 4 and 6 h.

**Figure 3:**
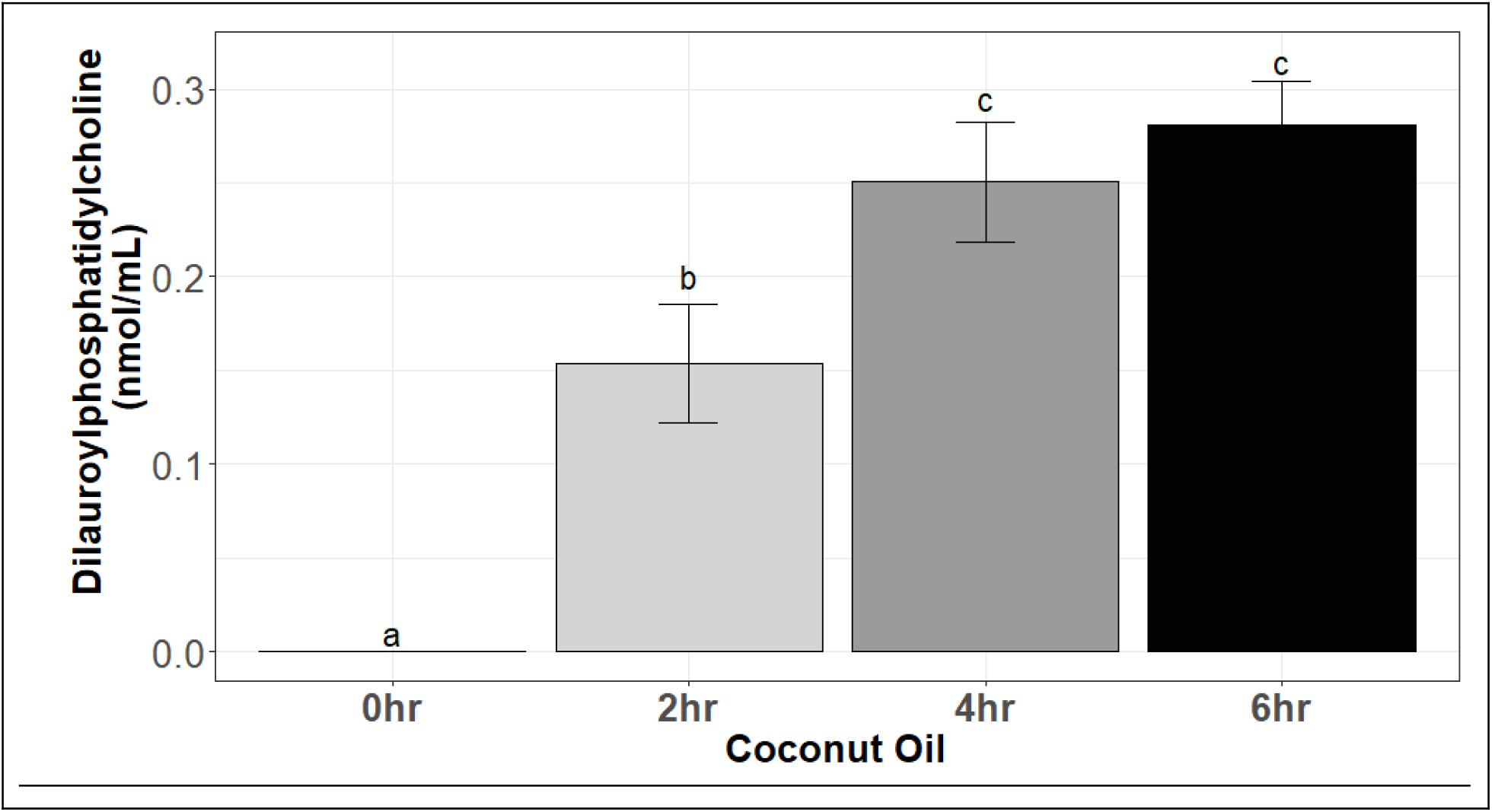
C12:0 Provision Results in the Appearance of DLPC in Postprandial Serum of Coconut Oil-Fed Healthy Human Subjects. Participants (n=10) completed a randomized, cross-over feeding trial consuming either coconut oil or olive oil-based breakfast smoothies assigned in random order. Following a 1-wk washout, participants consumed the opposite smoothie on a second visit. Baseline blood samples and postprandial blood samples were taken to assess the presence of DLPC in the plasma. Statistically significant (p<0.05) differences between time points were assessed by 1-Way ANOVA.

### C12:0 Provision Results in Induction of LRH-1 target gene expression *Ex Vivo* and *In Vivo*

While we observed the production of the LRH-1 ligand, DLPC, from the provision of its acyl constituent, C12:0, it remained unclear if this results in ligand-induced enhanced transcriptional activity. To address this, LRH-1^fl/fl^ mice and Albumin-Cre mice were bred to generate hepatic knockout (Cre+; LRH1-LKO) and wild-type (LRH1-WT; Cre-, LRH-1^fl/fl^ and LRH-1^fl/-^) and used to assess LRH-1 target gene expression in primary hepatocyte (ex vivo) and feeding (in vivo) experiments. Following treatment for 8 hours, primary hepatocytes (n=6 per treatment-x-genotype group) from LRH1-WT mice treated with C12:0 exhibited significantly increased expression of the LRH-1 target gene, CYP8B1, but not GNMT, relative to C16:0, C18:1n-9 or vehicle **(Figure 4 A,B)**; as expected, expression of both target genes was significantly reduced in LRH1-LKO mice relative to LRH1-WT, with no evidence of C12:0-induced enhanced transcriptional in LRH1-LKO mice. After 10 days of feeding, LRH1-WT mice fed C12:0 (n=7) exhibited significantly higher expression of both *CYP8B1* and *GNMT* relative to C16:0-fed LRH1-WT (n=7) as well as C12:0-(n=7) and C16:0-(n=9) fed LRH1-LKO mice. Collectively, these LRH-1 dependent, C12:0-induced increases in established LRH-1 target genes, are consistent with C12:0 provision resulting in functional LRH-1 ligand production.

**Figure 4:**
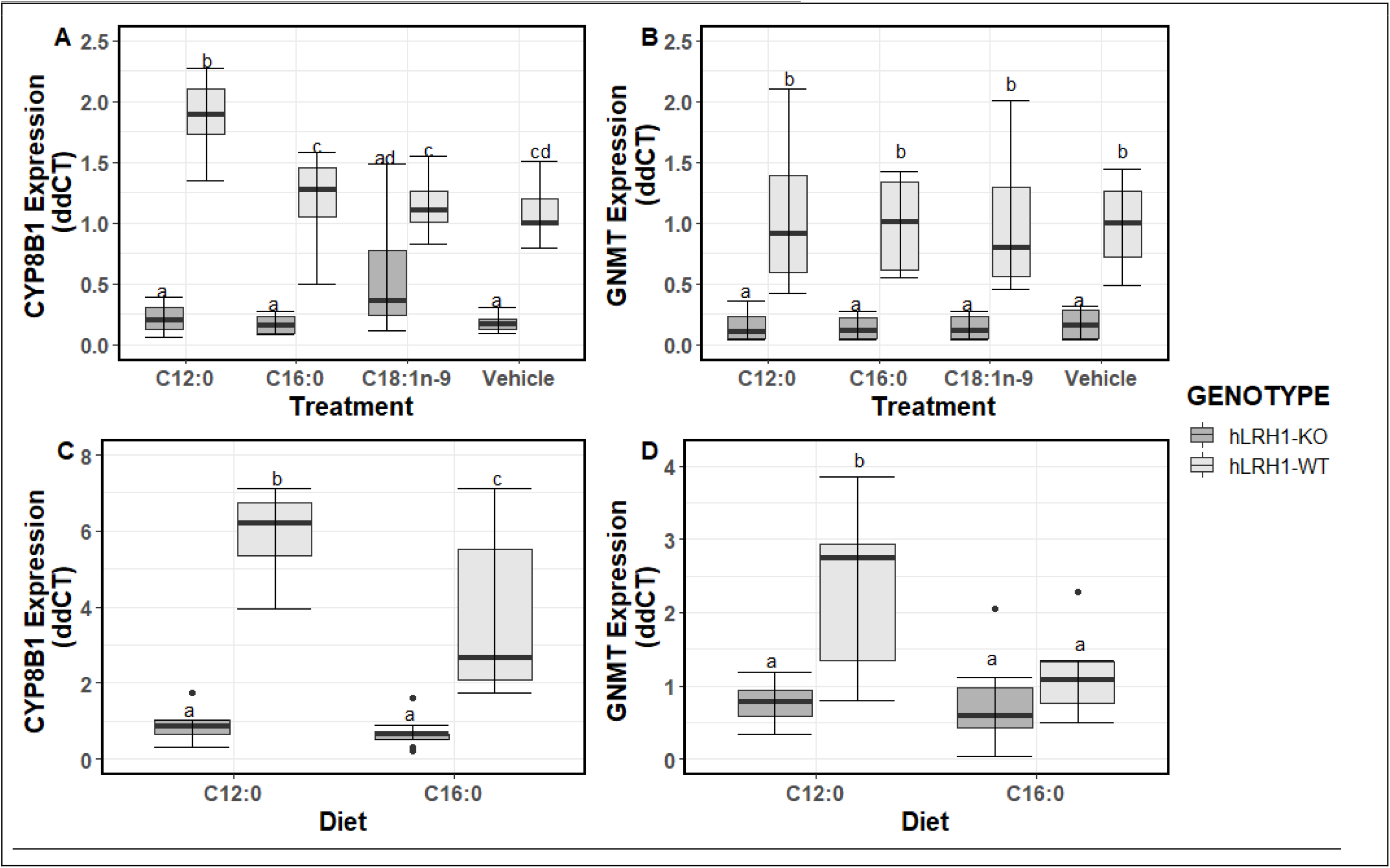
C12:0 Provision Results in LRH-1 Agonism Ex Vivo and In Vivo. A,B) Primary hepatocytes were isolated from hepatic LRH-1 wild type (hLRH1-WT) and hepatic LRH-1 knockout (hLRH-KO) mice (n=4; 2 per genotype) and underwent their indicated fatty acid or vehicle treatments for 8 hours. Expression of the LRH-1 target genes, CYP8B1 and GNMT, were assessed as evidence of LRH-1 activation. C,D) Hepatic LRH-1 wild type (hLRH1-WT) and hepatic LRH-1 knockout (hLRH-KO) mice were fed purified diets with 10% of kilocalories derived from C12:0 or C16:0 for 10 days; final sample sizes were as follows: hLRH1-WT-C12:0, n=7; hLRH1-WT-C16:0, n=7; hLRH1-KO-C12:0, n=7; hLRH1-KO-C16:0, n=9. Expression of the LRH-1 target genes, CYP8B1 and GNMT, were assessed as evidence of LRH-1 activation. Differences in gene expression were assessed using 2-way ANOVA with Tukey’s posthoc comparisons. Statistically significant differences between groups (p<0.05) are indicated via superscript lettering.

## DISCUSSION

Herein, we report the endogenous production of the LRH-1 agonist ligand DLPC following the provision of its acyl constituent, C12:0/lauric acid, across cultured cells, C57BL6J mice, and in a human randomized cross-over trial. Endogenous production of DLPC was not observed under conditions without the provision of exogenous C12:0/lauric acid, including under conditions of enhanced *de novo* lipogenesis in mice fed a 0% fat diet. We confirmed the need for exogenous C12:0 with ^13^C-labeled C12:0, and further demonstrated a reliance on the intestinal CDP-choline pathway of phosphatidylcholine biosynthesis to produce DLPC initially following intake, with evidence of complex whole body synthesis from plasma isotopic enrichment schemes. Provision of C12:0 both *ex vivo* and *in vivo* was associated with increased expression of hepatic LRH-1 target genes in wild-type, but not hepatic LRH-1 knockout mice, indicating that provision of C12:0 not only produces DLPC but results in LRH-1 activation.

Our investigation has strengths, limitations and need for follow-up investigation: We rigorously tested the hypothesis that the absence of DLPC is due to insufficient availability of its acyl constituent, C12:0, across multiple cell lines, acute and chronic animal feeding studies, and in a single-blind randomized controlled cross-over feeding trial. The production of DLPC by lauric feeding was complemented by the use of transgenic animals to show evidence of hepatic LRH-1 agonism via C12:0 supplementation *ex vivo* and feeding *in vivo*, an effect not observed in mice lacking hepatic LRH-1. Our investigation focused strongly on the effects of C12:0 provision relative to other purified fatty acids, whereas the literature/media commonly makes claims related to C12:0 from its classification as a medium chain fatty acid utilizing studies confounded by co-provision of additional, uniquely bioactive fatty acids^42^. Additionally, we incorporated stable isotope methodologies to assess inferences about the metabolic pathways required to produce DLPC, though the enzymatic steps required for DLPC production from C12:0 provision require further investigation.

While we have evidence of LRH-1 agonism following C12:0 provision, it is notable that there remains no way to specifically block DLPC synthesis following C12:0 provision, without additional confounding, to show that C12:0 does not influence LRH-1 through a non-DLPC pathway. It remains possible that other metabolites resulting from C12:0 provision may influence LRH-1 activity via ligand binding or other under-investigated mechanisms, such as the impact of C12:0 as a post-translational modification^19^. Though the amount of DLPC produced from C12:0 is small as a proportion of the total PC pool/per gram tissue, the subcellular localization of DLPC remains un-characterized. It is notable that PC pools differing in composition are compartmentalized in the cell, with endonuclear PC enriched in saturated fatty acids^43,44^. The functional purpose of this pool remains unclear, but may indeed be to facilitate local production of nuclear receptor ligands. Future investigations are necessary to determine whether C12:0 intakes result in preferential enrichment of C12:0-induced DLPC in endonuclear PC pools.

We set out to test whether C12:0 feeding resulted in LRH-1 agonism utilizing hepatocyte-specific LRH-1 knockout mice, due to the embryonic lethality of whole body LRH-1 knockout mice and the mild phenotype of heterozygous LRH-1 knockout mice. We utilized hepatocyte-specific LRH-1 knockout mice to test our hypothesis because of the well-defined hepatic LRH-1 cistrome^45^ and the role of LRH-1 in regulating target genes involved in multiple metabolic pathways (*CYP8B1*/lipid, *GNMT*/methyl) in this tissue. This unique characteristic of LRH-1 in the liver allowed us to limit the impact of *a priori* concerns unique to studies of nutrition, whereby the lack of a true placebo and need for choosing an isocaloric comparator introduces novel potential for confounding. Future studies investigating dietary-induced agonism of nuclear receptors on both transcriptional and physiological outcomes will require careful planning and interpretation of isocaloric macronutrient comparators, particularly in tissues with few known LRH-1 targets that are metabolically independent of one another. For example, C16:0 may or may not be an appropriate comparator for intestine-specific LRH-1 knockout studies, whereby inflammatory triggers are key to inducing LRH-1 transcriptional networks and influencing CYP11A1 expression^46^.

Lastly, our feeding study, while highly controlled and utilizing a strong cross-over design, does not show evidence for LRH-1 agonism; future studies with C12:0 feeding and tissue biopsies will be required to assess this. Our study notably needs to be repeated in disease-specific populations to ensure uncompromised DLPC production/presence in target tissues, as well as in longer term feeding trials with diverse metabolic, surrogate and quality of life outcomes. Certain disease states where LRH-1 is clearly implicated, such as inflammatory bowel disorders^46–48^, are high priority for investigating C12:0-modified diets and medical nutrition products.

There are several implications of our work, for both the fields of nuclear receptor biology and nutritional sciences. Our results highlight the importance of considering the fatty acid composition of nutrient exposures when assessing potential ligands for nuclear receptors. Indeed, the lack of C12:0 in culture mediums and laboratory animal diets likely facilitated the identification of DLPC as a LRH-1 ligand, and suggest that manipulation of fatty acid availability may reveal other potential agonists relevant to nuclear receptor signaling.

Our study findings, coupled with several observations throughout the literature, contribute to ongoing controversies over the impact of saturated fatty acids on human health. Current dietary recommendations state that food sources of saturated fatty acids should be limited, and recent recommendations have specifically highlighted the potential negative effects of coconut oil^49,50^. The World Health Organization specifically recommends reductions in the intake of three common hypercholesteremic saturated fatty acids: C12:0 (lauric acid), C14:0 (myristic acid) and C16:0 (palmitic acid)^51^. All three of these fatty acids raise LDL-C^20^, a causal factor in the development of cardiovascular disease^52^. To date, randomized controlled trials testing the hypothesis that cholesterol lowering diets, of which reduced saturated fat consumption are a core component, have primarily limited fat sources abundant in C14:0 and C16:0 (e.g. high fat meats and dairy products)^53^. No trial has tested the impact of C12:0 rich oils (e.g. coconut, palm kernel oil) on disease endpoints, and thus, decisions about health impacts of these are limited to surrogate endpoints. Assessing the health impact of an intervention via a single surrogate endpoint has significant limitations^54–56^, particularly in that pleiotropic effects of interventions may modify the overall clinical effect size on actual disease endpoints. In the case of nutrients, pleiotropic effects of nutrients are often context-dependent, and may over- or under-estimate the impacts on health outcomes.

There is evidence that C12:0 may have effects quite distinct from those of other saturated fatty acids consistent with pleitropy. Notably, coconut oil and/or lauric have been consistently observed as outlier sources of saturated fat in the preclinical and clinical literature assessing cardiometabolic outcomes ^21,25,27–30,57–60^. Since the earliest investigations by Paigen and colleagues defining atherogenic diets for use in murine models, saturated fatty acids have been shown to be highly correlated with atherosclerosis lesion size - with coconut oil being the notable exception^25^. Coconut oil and C12:0 also failed to induce insulin resistance to the degree that various comparator oils/fats do in rodent models^27,57,58^. Protective cardiometabolic effects of C12:0 in human feeding trials^28^ and epidemiological ^30 29^ investigations, relative to longer-chain saturated fats, have also begun to emerge. Pleiotropic effects of C12:0 through LRH-1 signaling, which has well-described roles in glucose sensing and whole body cholesterol regulation, protecting against insulin resistance^61^, beta cell dysfunction^62,63^, and atherosclerosis^64^, remains speculative but plausible mechanism by which C12:0, and by extension, C12:0-rich oils such as palm kernel oil and coconut oil, may have diverse impacts on cardiometabolic health. Advancing speculation and ultimately, our understanding of C12:0 on health outcomes will require tissue-specific-LRH-1 knockout murine feeding trials, well-designed human feeding trials with multiple isocaloric comparators and surrogate endpoints across a range of intakes, and a revisiting of dietary assessment methodologies in large epidemiological studies to capture lauric-acid containing oils. While much of this discussion surrounds cardiometabolic health, the impact of C12:0 intake (increased or reduced) on other disease categories influenced by LRH-1 are worthwhile, including pathophysiological states characterized by reduced ovarian progesterone synthesis (e.g. recurrent pregnancy loss)^65,66^, cancer^67,68^ and intestinal inflammatory conditions^46^. Additionally, the high content of C12:0 in mammalian milk warrants investigation into the potential relevance of LRH-1 agonism at this life-stage ^23^.

Ultimately, substantial research on the relevance of DLPC-induced LRH-1 agonism in humans, either by diet or pharmacological administration, is essential before dietary recommendations about C12:0 should be altered.

## CONCLUSIONS

Herein, we demonstrate the endogenous production of DLPC, an agonist ligand for the orphan nuclear receptor LRH-1, following the provision of the saturated fatty acid, C12:0/lauric acid. C12:0 provision increases expression of LRH-1 target genes, as expected from the agonist activity of DLPC, an effect that is lost in the genetic absence of LRH-1. These results support the potential for pleiotropic effects of C12:0 and warrant further investigation into the physiological and pathophysiological effects of C12:0-containing oils.

## REFERENCES

1. Sever, R. & Glass, C. K. Signaling by nuclear receptors. Cold Spring Harb. Perspect. Biol. 5, a016709 (2013).

2. Fayard, E., Auwerx, J. & Schoonjans, K. LRH-1: an orphan nuclear receptor involved in development, metabolism and steroidogenesis. Trends Cell Biol. 14, 250–260 (2004).

3. Tissue expression of MFSD2A - Summary - The Human Protein Atlas. at <https://www.proteinatlas.org/ENSG00000168389-MFSD2A/tissue>

4. Bookout, A. L. et al. Anatomical profiling of nuclear receptor expression reveals a hierarchical transcriptional network. Cell 126, 789–799 (2006).

5. Wagner, M. et al. Liver receptor homolog-1 is a critical determinant of methyl-pool metabolism. Hepatology 63, 95–106 (2016).

6. Forman, B. M. Are those phospholipids in your pocket? Cell Metab. 1, 153–155 (2005).

7. Sablin, E. P. et al. Structure of Liver Receptor Homolog-1 (NR5A2) with PIP3 hormone bound in the ligand binding pocket. J. Struct. Biol. 192, 342–348 (2015).

8. Ortlund, E. A. et al. Modulation of human nuclear receptor LRH-1 activity by phospholipids and SHP. Nat. Struct. Mol. Biol. 12, 357–363 (2005).

9. Wang, W. et al. The crystal structures of human steroidogenic factor-1 and liver receptor homologue-1. Proc Natl Acad Sci USA 102, 7505–7510 (2005).

10. Lee, J. M. et al. A nuclear-receptor-dependent phosphatidylcholine pathway with antidiabetic effects. Nature 474, 506–510 (2011).

11. Lefèvre, L. et al. LRH-1 mediates anti-inflammatory and antifungal phenotype of IL-13- activated macrophages through the PPARγ ligand synthesis. Nat. Commun. 6, 6801 (2015).

12. Choi, S. et al. Methyl-Sensing Nuclear Receptor Liver Receptor Homolog-1 Regulates Mitochondrial Function in Mouse Hepatocytes. Hepatology 71, 1055–1069 (2020).

13. Bolado-Carrancio, A., Riancho, J. A., Sainz, J. & Rodríguez-Rey, J. C. Activation of nuclear receptor NR5A2 increases Glut4 expression and glucose metabolism in muscle cells. Biochem. Biophys. Res. Commun. 446, 614–619 (2014).

14. Human Metabolome Database: Search Results for metabolite. at <http://www.hmdb.ca/unearth/q?utf8=%E2%9C%93&query=PC%2812%3A0%2F12%3A0%29&searcher=metabolites&button=>

15. mouse Multiple tissue Metabolome DataBase. at <http://mmdb.iab.keio.ac.jp/search.html>

16. Hellerstein, M. K. De novo lipogenesis in humans: metabolic and regulatory aspects. Eur. J. Clin. Nutr. 53 Suppl 1, S53–65 (1999).

17. Klatt, K. & Caudill, M. in Biochemical, Molecular and Physiological Aspects of Human Nutrition (eds. Stipanuk, M. & Caudill, M.) (Elsevier).

18. Oliveira, M. M. & Vaughan, M. Incorporation of fatty acids into phospholipids of erythrocyte membranes. J. Lipid Res. 5, 156–162 (1964).

19. Rioux, V., Daval, S., Guillou, H., Jan, S. & Legrand, P. Although it is rapidly metabolized in cultured rat hepatocytes, lauric acid is used for protein acylation. Reprod. Nutr. Dev. 43, 419–430 (2003).

20. WHO | Effects of saturated fatty acids on serum lipids and lipoproteins: a systematic review and regression analysis. at <http://www.who.int/nutrition/publications/nutrientrequirements/sfa_systematic_review/en/>

21. Prior, I. A., Davidson, F., Salmond, C. E. & Czochanska, Z. Cholesterol, coconuts, and diet on Polynesian atolls: a natural experiment: the Pukapuka and Tokelau island studies. Am. J. Clin. Nutr. 34, 1552–1561 (1981).

22. Food Composition Databases Show Nutrients List. at <https://ndb.nal.usda.gov/ndb/nutrients/index>

23. Mosley, E. E., Wright, A. L., McGuire, M. K. & McGuire, M. A. trans Fatty acids in milk produced by women in the United States. Am. J. Clin. Nutr. 82, 1292–1297 (2005).

24. Mosby, J. M., Wodzicki, K. & Shorland, F. B. Fatty acid composition of the depot fats of the Polynesian rat, *Rattus exulans*, Tokelau Islands. N.Z. J. Zool. 1, 67–70 (1974).

25. Nishina, P. M. et al. Effects of dietary fats from animal and plant sources on diet-induced fatty streak lesions in C57BL/6J mice. J. Lipid Res. 34, 1413–1422 (1993).

26. Merkel, M., Velez-Carrasco, W., Hudgins, L. C. & Breslow, J. L. Compared with saturated fatty acids, dietary monounsaturated fatty acids and carbohydrates increase atherosclerosis and VLDL cholesterol levels in LDL receptor-deficient, but not apolipoprotein E-deficient, mice. Proc Natl Acad Sci USA 98, 13294–13299 (2001).

27. Buettner, R. et al. Defining high-fat-diet rat models: metabolic and molecular effects of different fat types. J. Mol. Endocrinol. 36, 485–501 (2006).

28. Lundsgaard, A.-M. et al. Small Amounts of Dietary Medium-Chain Fatty Acids Protect Against Insulin Resistance During Caloric Excess in Humans. Diabetes (2020). doi:10.2337/db20-0582

29. Praagman, J. et al. Consumption of individual saturated fatty acids and the risk of myocardial infarction in a UK and a Danish cohort. Int. J. Cardiol. 279, 18–26 (2019).

30. Liu, S., van der Schouw, Y. T., Soedamah-Muthu, S. S., Spijkerman, A. M. W. & Sluijs, I. Intake of dietary saturated fatty acids and risk of type 2 diabetes in the European Prospective Investigation into Cancer and Nutrition-Netherlands cohort: associations by types, sources of fatty acids and substitution by macronutrients. Eur. J. Nutr. 58, 1125–1136 (2019).

31. DeLany, J. P., Windhauser, M. M., Champagne, C. M. & Bray, G. A. Differential oxidation of individual dietary fatty acids in humans. Am. J. Clin. Nutr. 72, 905–911 (2000).

32. Lee, J. Y. et al. Reciprocal modulation of Toll-like receptor-4 signaling pathways involving MyD88 and phosphatidylinositol 3-kinase/AKT by saturated and polyunsaturated fatty acids. J. Biol. Chem. 278, 37041–37051 (2003).

33. Lee, J. Y. et al. Saturated fatty acid activates but polyunsaturated fatty acid inhibits Toll-like receptor 2 dimerized with Toll-like receptor 6 or 1. J. Biol. Chem. 279, 16971–16979 (2004).

34. Lee, J. Y., Sohn, K. H., Rhee, S. H. & Hwang, D. Saturated fatty acids, but not unsaturated fatty acids, induce the expression of cyclooxygenase-2 mediated through Toll-like receptor 4. J. Biol. Chem. 276, 16683–16689 (2001).

35. Mamrosh, J. L. et al. Nuclear receptor LRH-1/NR5A2 is required and targetable for liver endoplasmic reticulum stress resolution. elife 3, e01694 (2014).

36. Chakravarthy, M. V. et al. Identification of a physiologically relevant endogenous ligand for PPARalpha in liver. Cell 138, 476–488 (2009).

37. Wei, Y. et al. MRG15 orchestrates rhythmic epigenomic remodelling and controls hepatic lipid metabolism. Nat. Metab. 2, 447–460 (2020).

38. Koc, H., Mar, M.-H., Ranasinghe, A., Swenberg, J. A. & Zeisel, S. H. Quantitation of choline and its metabolites in tissues and foods by liquid chromatography/electrospray ionization-isotope dilution mass spectrometry. Anal. Chem. 74, 4734–4740 (2002).

39. Yan, J. et al. Pregnancy alters choline dynamics: results of a randomized trial using stable isotope methodology in pregnant and nonpregnant women. Am. J. Clin. Nutr. 98, 1459–1467 (2013).

40. Pynn, C. J. et al. Specificity and rate of human and mouse liver and plasma phosphatidylcholine synthesis analyzed in vivo. J. Lipid Res. 52, 399–407 (2011).

41. Shields, D. J., Agellon, L. B. & Vance, D. E. Structure, expression profile and alternative processing of the human phosphatidylethanolamine N-methyltransferase (PEMT) gene. Biochim. Biophys. Acta 1532, 105–114 (2001).

42. Eyres, L., Eyres, M. F., Chisholm, A. & Brown, R. C. Coconut oil consumption and cardiovascular risk factors in humans. Nutr. Rev. 74, 267–280 (2016).

43. Hunt, A. N., Clark, G. T., Attard, G. S. & Postle, A. D. Highly saturated endonuclear phosphatidylcholine is synthesized in situ and colocated with CDP-choline pathway enzymes. J. Biol. Chem. 276, 8492–8499 (2001).

44. Hunt, A. N., Clark, G. T., Neale, J. R. & Postle, A. D. A comparison of the molecular specificities of whole cell and endonuclear phosphatidylcholine synthesis. FEBS Lett. 530, 89–93 (2002).

45. Chong, H. K., Biesinger, J., Seo, Y.-K., Xie, X. & Osborne, T. F. Genome-wide analysis of hepatic LRH-1 reveals a promoter binding preference and suggests a role in regulating genes of lipid metabolism in concert with FXR. BMC Genomics 13, 51 (2012).

46. Coste, A. et al. LRH-1-mediated glucocorticoid synthesis in enterocytes protects against inflammatory bowel disease. Proc Natl Acad Sci USA 104, 13098–13103 (2007).

47. Liu, J. Z. et al. Association analyses identify 38 susceptibility loci for inflammatory bowel disease and highlight shared genetic risk across populations. Nat. Genet. 47, 979–986 (2015).

48. Bayrer, J. R. et al. LRH-1 mitigates intestinal inflammatory disease by maintaining epithelial homeostasis and cell survival. Nat. Commun. 9, 4055 (2018).

49. 2015-2020 Dietary Guidelines - health.gov. at <https://health.gov/dietaryguidelines/2015/guidelines/>

50. Sacks, F. M. Coconut oil and heart health. Circulation 141, 815–817 (2020).

51. WHO | 5. Population nutrient intake goals for preventing diet-related chronic diseases. at <http://www.who.int/nutrition/topics/5_population_nutrient/en/index10.html>

52. Ference, B. A. et al. Low-density lipoproteins cause atherosclerotic cardiovascular disease. 1. Evidence from genetic, epidemiologic, and clinical studies. A consensus statement from the European Atherosclerosis Society Consensus Panel. Eur. Heart J. 38, 2459–2472 (2017).

53. Hooper, L., Martin, N., Abdelhamid, A. & Davey Smith, G. Reduction in saturated fat intake for cardiovascular disease. Cochrane Database Syst. Rev. CD011737 (2015). doi:10.1002/14651858.CD011737

54. Bikdeli, B. et al. Two Decades of Cardiovascular Trials With Primary Surrogate Endpoints: 1990-2011. J. Am. Heart Assoc. 6, (2017).

55. Tall, A. R. & Rader, D. J. Trials and tribulations of CETP inhibitors. Circ. Res. 122, 106–112 (2018).

56. Institute of Medicine (US) Committee on Qualification of Biomarkers and Surrogate Endpoints in Chronic Disease. Evaluation of biomarkers and surrogate endpoints in chronic disease. (National Academies Press (US), 2010). doi:10.17226/12869

57. Saraswathi, V. et al. Lauric Acid versus Palmitic Acid: Effects on Adipose Tissue Inflammation, Insulin Resistance, and Non-Alcoholic Fatty Liver Disease in Obesity. Biology (Basel) 9, (2020).

58. Deol, P. et al. Soybean Oil Is More Obesogenic and Diabetogenic than Coconut Oil and Fructose in Mouse: Potential Role for the Liver. PLoS ONE 10, e0132672 (2015).

59. Xia, J. et al. Lauric Triglyceride Ameliorates High-Fat-Diet-Induced Obesity in Rats by Reducing Lipogenesis and Increasing Lipolysis and β-Oxidation. J. Agric. Food Chem. (2021). doi:10.1021/acs.jafc.0c07342

60. Žáček, P. et al. Dietary saturated fatty acid type impacts obesity-induced metabolic dysfunction and plasma lipidomic signatures in mice. J. Nutr. Biochem. 64, 32–44 (2019).

61. Oosterveer, M. H. et al. LRH-1-dependent glucose sensing determines intermediary metabolism in liver. J. Clin. Invest. 122, 2817–2826 (2012).

62. Baquié, M. et al. The liver receptor homolog-1 (LRH-1) is expressed in human islets and protects {beta}-cells against stress-induced apoptosis. Hum. Mol. Genet. 20, 2823–2833 (2011).

63. Cobo-Vuilleumier, N. et al. LRH-1 agonism favours an immune-islet dialogue which protects against diabetes mellitus. Nat. Commun. 9, 1488 (2018).

64. Stein, S. et al. SUMOylation-dependent LRH-1/PROX1 interaction promotes atherosclerosis by decreasing hepatic reverse cholesterol transport. Cell Metab. 20, 603–613 (2014).

65. Duggavathi, R. et al. Liver receptor homolog 1 is essential for ovulation. Genes Dev. 22, 1871–1876 (2008).

66. Zhang, C. et al. Liver receptor homolog-1 is essential for pregnancy. Nat. Med. 19, 1061–1066 (2013).

67. Kramer, H. B. et al. LRH-1 drives colon cancer cell growth by repressing the expression of the CDKN1A gene in a p53-dependent manner. Nucleic Acids Res. 44, 582–594 (2016).

68. Bianco, S., Jangal, M., Garneau, D. & Gévry, N. LRH-1 controls proliferation in breast tumor cells by regulating CDKN1A gene expression. Oncogene 34, 4509–4518 (2015).

