## Supplementary Figures and Tables for "LRH-1 NUTRIGENOMICS: The Provision of Lauric Acid Results in the Endogenous Production of the Liver Receptor Homolog-1 Ligand, Dilauroylphosphatidylcholine, and LRH-1 Transactivation"

**Supplementary Figure 1: NHANES C12:0 Intakes**

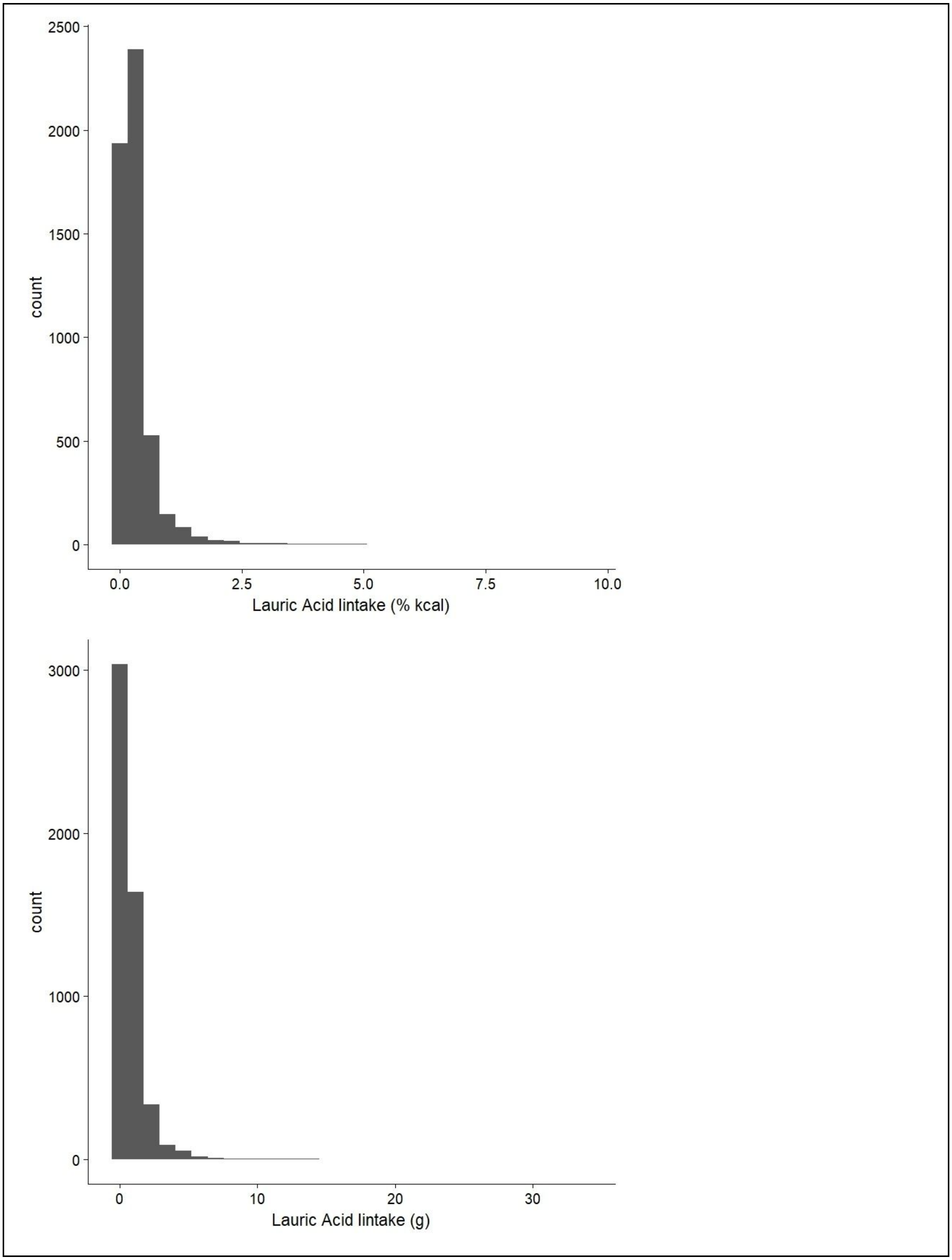

**Supplementary Figure 2: CONSORT Flow Diagram**

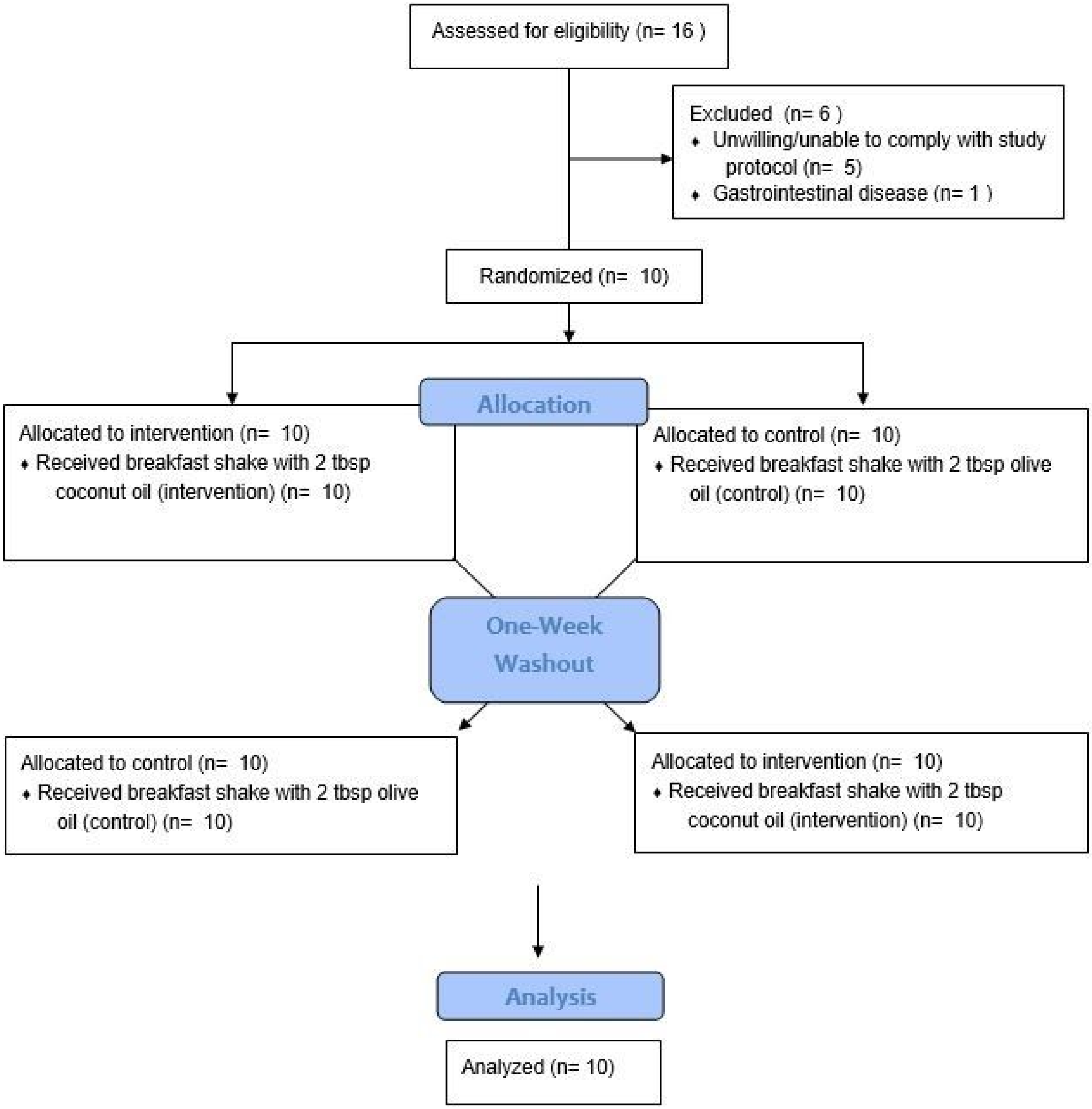

**Supplementary Figure 3: Purified Diet Composition**

| Product # | D20061101 | | D20061102 | |
|---|---|---|---|---|
| % | gm | *kcal* | gm | *kcal* |
| Protein | 20 | *20* | 20 | *20* |
| Carbohydrate | 60 | *59* | 60 | *59* |
| Fat | 10 | *21* | 10 | *21* |
| Total | | *100* | | *100* |
| kcal/gm | 4.1 | | 4.1 | |
| **Ingredient** | **gm** | ***kcal*** | **gm** | ***kcal*** |
| Casein | 200 | *800* | 200 | *800* |
| L-Cystine | 3 | *12* | 3 | *12* |
| Corn Starch | **314.5** | ***1258*** | **314.5** | ***1258*** |
| Maltodextrin 10 | 100 | *400* | 100 | *400* |
| Sucrose | 172.8 | *691* | 172.8 | *691* |
| Cellulose, BW200 | 50 | *0* | 50 | *0* |
| Soybean Oil | 25 | *225* | 25 | *225* |
| Lard | 25.00 | *225* | 25.00 | *225* |
| Palmitic Acid | **45.08** | ***406*** | 0 | *0* |
| Lauric Acid | 0 | *0* | **45.08** | ***406*** |
| Oleic Acid | 0 | *0* | 0 | *0* |
| Mineral Mix S10026 | 10 | *0* | 10 | *0* |
| DiCalcium Phosphate | 13 | *0* | 13 | *0* |
| Calcium Carbonate | 5.5 | *0* | 5.5 | *0* |
| Potassium Citrate, 1 H2O | 16.5 | *0* | 16.5 | *0* |
| Vitamin Mix V10001 | 10 | *40* | 10 | *40* |
| Choline Bitartrate | 2 | *0* | 2 | *0* |
| FD&C Yellow Dye #5 | 0 | *0* | 0.05 | *0* |
| FD&C Red Dye #40 | 0 | *0* | 0 | *0* |
| FD&C Blue Dye #1 | 0.05 | *0* | 0 | *0* |
| **Total** | **992.43** | ***4057*** | **992.43** | ***4057*** |
| Palmitic Acid (%) | 10.00 | | | |
| Lauric Acid (%) | | | 10.00 | |

**Supplementary Table 1: High C12:0 Feeding Recipe**

| High C12:0 Diet | g/kg of diet |
|---|---|
| Added Fat | 250 |
| Mineral | 35 |
| Vitamin | 10 |
| 2018 Teklad Chow | 705 |

**Supplementary Table 2: Trial Participant Descriptives**

| ID | Sex | Age | Weight (kg) | Height (cm) | BMI |
|---|---|---|---|---|---|
| L1 | F | 28 | 61.2 | 170 | 21.2 |
| L2 | M | 26 | 81.6 | 180 | 25.2 |
| L3 | F | 33 | 63.5 | 175.5 | 20.6 |
| L4 | F | 46 | 65.7 | 170 | 22.7 |
| L5 | F | 23 | 56.5 | 173 | 18.9 |
| L6 | F | 24 | 63.5 | 162.5 | 24 |
| L7 | F | 27 | 58 | 162.5 | 22 |
| L8 | F | 28 | 57 | 172 | 19.3 |
| L9 | F | 27 | 68 | 167.5 | 24.2 |
| L10 | M | 30 | 99.85 | 190.5 | 27.5 |

**Supplementary Table 3: Mouse Primer Sequences**

| ***Primer*** | ***Direction*** | ***Sequence*** *(5' -> 3')* |
|---|---|---|
| ***mGAPDH*** | *Forward Primer* | *CTTTGGCATTGTGGAAGGGC* |
| | *Reverse Primer* | *CAGGGATGATGTTCTGGGCA* |
| ***mCYP8B1*** | *Forward Primer* | *TTGCAAATGCTGCCTCAACC* |
| | *Reverse Primer* | *TAACAGTCGCACACATGGCT* |
| ***mGNMT*** | *Forward Primer* | GTGCTGGACGTAGCCTGT |
| | *Reverse Primer* | ATCACGCTGAAGCCCTCTT |
